# Splatter: simulation of single-cell RNA sequencing data

**DOI:** 10.1101/133173

**Authors:** Luke Zappia, Belinda Phipson, Alicia Oshlack

**Affiliations:** Murdoch Childrens Research Institute, Royal Children's Hospital, 50 Flemington Rd, Parkville VIC 3052, Australia; School of Biosciences, The University of Melbourne, Parkville VIC 3010, Australia

**Keywords:** Single-cell, RNA-seq, Simulation, Software

## Abstract

As single-cell RNA sequencing technologies have rapidly developed, so have analysis methods. Many methods have been tested, developed and validated using simulated datasets. Unfortunately, current simulations are often poorly documented, their similarity to real data is not demonstrated, or reproducible code is not available.

Here we present the Splatter Bioconductor package for simple, reproducible and well-documented simulation of single-cell RNA-seq data. Splatter provides an interface to multiple simulation methods including Splat, our own simulation, based on a gamma-Poisson distribution. Splat can simulate single populations of cells, populations with multiple cell types or differentiation paths.

## Background

The first decade of next-generation sequencing has seen an explosion in our understanding of the genome [1]. In particular, the development of RNA sequencing (RNA-seq) has enabled unprecedented insight into the dynamics of gene expression [2]. Researchers now routinely conduct experiments designed to test how gene expression is affected by various stimuli. One limitation of bulk RNA-seq experiments is that they measure the average expression level of genes across the many cells in a sample. However, recent technological developments have enabled the extraction and amplification of minute quantities of RNA, allowing sequencing to be conducted on the level of single cells [3]. The increased resolution of single-cell RNA-seq (scRNA-seq) data has made a range of new analyses possible.

As scRNA-seq data has become available there has been a rapid development of new bioinformatics tools attempting to unlock its potential. Currently there are at least 120 software packages that have been designed specifically for the analysis of scRNA-seq data, the majority of which have been published in peer-reviewed journals or as preprints [4]. The focus of these tools is often different from those designed for the analysis of a bulk RNA-seq experiment. In a bulk experiment, the groups of samples are known and a common task is to test for genes that are differentially expressed (DE) between two or more groups. In contrast, the groups in a single-cell experiment are usually unknown and the analysis is often more exploratory.

Much of the existing software focuses on assigning cells to groups based on their expression profiles (clustering) before applying more traditional DE testing. This approach is taken by tools such as SC3 [5], CIDR [6] and Seurat [7], and is appropriate for a sample of mature cells where it is reasonable to expect cells to have a particular type. In a developmental setting, for example, where stem cells are differentiating into mature cells, it may be more appropriate to order cells along a continuous trajectory from one cell type to another. Tools such as Monocle [8], CellTree [9] and Sincell [10] take this approach, ordering cells along a path, then identifying patterns in the changes of gene expression along that path.

Another defining characteristic of scRNA-seq data is its sparsity; typically expression is only observed for relatively few genes in each cell. The observed zero counts have both biological (different cell types express different genes) and technical (an expressed RNA molecule might not be captured) causes, with technical zeros often referred to as “dropout”. Some analysis methods (ZIFA [11], MAST [12], ZINB-WaVE [13]) incorporate dropout into their models while others (MAGIC [14], SAVER [15], scImpute [16]) attempt to infer what the true expression levels should be.

Existing scRNA-seq analysis packages, and any new methods that are being developed, should demonstrate two properties: first that they can do what they claim to do, whether that is clustering, lineage tracing, differential expression testing or improved performance compared to other methods, and second that they produce some meaningful biological insight. The second criterion is specific to particular studies but it should be possible to address the first point in a more general way.

A common way to test the performance of an analysis method is through a simulation. Simulated data provides a known truth to test against, making it possible to assess whether a method has been implemented correctly, whether the assumptions of the method are appropriate, and demonstrating the method's limitations. Such tests are often difficult with real biological data, as an experiment must be specifically designed, or results from an appropriate orthogonal test taken as the truth. Simulations, however, easily allow access to a range of metrics for assessing the performance of an analysis method. An additional advantage of evaluating methods using simulated data is that many datasets, with different parameters and assumptions, can be rapidly generated at minimal cost. As such, many of the scRNA-seq analysis packages that are currently available have used simulations to demonstrate their effectiveness. These simulations, however, are often not described in a reproducible or reusable way and the code to construct them may not be readily available. When code is available it may be poorly documented or written specifically for the computing environment used by the authors, limiting its reproducibility and making it difficult for other researchers to reuse. Most importantly, publications do not usually provide sufficient detail demonstrating that a simulation is similar to real datasets, or in what ways it differs.

In this paper we present Splatter, an R Bioconductor package for reproducible and accurate simulation of single-cell RNA sequencing data. Splatter is a framework designed to provide a consistent interface to multiple published simulations, enabling researchers to quickly simulate scRNA-seq count data in a reproducible fashion and make comparisons between simulations and real data. Along with the framework we have developed our own simulation model, Splat, and show how it compares to previously published simulations based on real datasets. We also provide a short example of how simulations can be used for assessing analysis methods.

## Results

### The Splatter framework

Currently, Splatter implements six different simulation models, each with their own assumptions but accessed through a consistent, easy-to-use interface. These simulations are described in more detail in the following sections and in the documentation for each simulation in Splatter, which also describes the required input parameters.

The Splatter simulation process consists of two steps. The first step estimates the parameters required for the simulation from a real dataset. The result of the first step is a parameters object unique to each simulation model. These objects have been designed to hold the information required for the specific simulation and display details such as which parameters can be estimated and which have been changed from the default value. It is important that each simulation has its own object for storing parameters as different simulations can vary greatly in the information they require. For example, some simulations only need parameters for well-known statistical distributions while others require large vectors or matrices of data sampled from real datasets.

In the second step, Splatter uses the estimated parameters, along with any additional parameters that cannot be estimated or are overridden by the user, to generate a synthetic scRNA-seq dataset. If there is no relevant real data to estimate parameters from, a synthetic dataset can still be generated using default parameters that can be manually modified by the user. Additional parameters that may be required depend on the simulation; these could include parameters indicating whether to use a zero-inflated model or the number of genes and cells to simulate. The main result of the simulation step is a matrix of counts, which is returned as an SCESet object as defined by the scater package. Scater is a low-level analysis package that provides various functions for quality control, visualization and preprocessing of scRNA-seq data [17]. Briefly, the structure of the SCESet combines cell by feature (gene) matrices for storing expression values along with tables for storing metadata about cells and features (further details are described in the scater documentation and the accompanying paper). This is a convenient format for returning intermediate values created during simulation as well as the final expression matrix. For example, the underlying gene expression means in different groups of cells are returned and could be used as a truth when evaluating differential expression testing. Using an SCESet also provides easy access to scater's functionality.

Splatter is also able to compare SCESet objects. These may contain simulations with different models or different parameters, or real datasets from which parameters have been estimated. The comparison function takes one or more SCESet objects, combines them (keeping any cell or gene-level information that is present in all of them) and produces a series of diagnostic plots comparing aspects of scRNA-seq data. The combined datasets are also returned, making it easy to produce additional comparison plots or statistics. Alternatively, one SCESet can be designated as a reference, such as the real data used to estimate parameters, and the difference between the reference and the other datasets can be assessed. This approach is particularly useful for comparing how well simulations recapitulate real datasets. Examples of these comparison plots are shown in the following sections.

### Simulation models

Splatter provides implementations of our own simulation model, Splat, as well as several previously published simulations. The previous simulations have either been published as R code associated with a paper or as functions in existing packages. By including them in Splatter, we have made them available in a single place in a more accessible way. If only a script was originally published, such as the Lun [18] and Lun 2 [19] simulations, the simulations have been re-implemented in Splatter. If the simulation is available in an existing R package, for example scDD [20] and BASiCS [21], we have simply written wrappers that provide consistent input and output but use the package implementation. We have endeavored to keep the simulations and estimation procedures as close as possible to what was originally published while providing a consistent interface within Splatter. The six different simulations currently available in Splatter are described below.

### Simple

The negative binomial is the most common distribution used to model RNA-seq count data, as in the edgeR [22] and DESeq [23] packages. The Simple simulation is a basic implementation of this approach. A mean expression level for each gene is simulated using a gamma distribution and the negative binomial distribution is used to generate a count for each cell based on these means, with a fixed dispersion parameter (default = 0.1) (Additional File 1 Figure 1). This simulation is primarily included as a baseline reference and is not intended to accurately reproduce many of the features of scRNA-seq data.

### Lun

Published in “Pooling across cells to normalize single-cell RNA sequencing data with many zero counts” [18] the Lun simulation builds on the Simple simulation by adding a scaling factor for each cell (Additional File 1 Figure 2). The cell factors are randomly sampled from a normal distribution with mean 1 and variance 0.5. The inverse-log_2_ transformed factors are used to adjust the gene means resulting in a matrix in which each cell has a different mean. This represents the kinds of technical effects that scaling normalisation aims to remove. The matrix of means is then used to sample counts from a negative binomial distribution, with a fixed dispersion parameter. This simulation can also model differential expression between multiple groups with fixed fold changes.

### Lun 2

In “Overcoming confounding plate effects in differential expression analyses of single-cell RNA-seq data” [19] Lun and Marioni extended the negative binomial model from the Lun simulation. This simulation samples input parameters from real data, with very little random sampling from statistical distributions. In the Lun 2 simulation the cell factors are replaced with a library size factor and an additional level of variation is added by including a batch effects factor. While the library size factor acts on individual cells the batch effects are applied to groups of cells from the same batch. This simulation is thus highly specific to the scenario when there are known batch effects present in the data, for example Fluidigm C1 plate effects. Differential expression can be added between two sets of batches and the user can choose to use a zero-inflated negative binomial (ZINB) model. Counts are simulated from a negative binomial using the library size and plate factor adjusted gene means and gene-wise dispersion estimates obtained from the real data. If the ZINB model is chosen, zero inflated estimates of gene means and dispersions are used instead. An additional step then randomly sets some counts to zero, based on the gene-wise proportions of zeros observed in the data. Additional File 1 Figure 3 shows the model assumptions and parameters for this simulation.

### scDD

The scDD package aims to test for differential expression between two groups of cells but also more complex changes such as differential distributions or differential proportions [20]. This is reflected in the scDD simulation, which can contain a mixture of genes simulated to have different distributions, or differing proportions where the expression of the gene is multi-modal. This simulation also samples information from a real dataset. As the scDD simulation is designed to reproduce a high quality, filtered dataset, it only samples from genes with less than 75 percent zeros. As a result, it only simulates relatively highly expressed genes. The Splatter package simply provides wrapper functions to the simulation function in the scDD package, while capturing the necessary inputs and outputs needed to compare to other simulations. The full details of the scDD simulation are described in the scDD package vignette [24].

### BASiCS

The BASiCS package introduced a model for separating variation in scRNA-seq data into biological and technical components based on the expression of external spike-in controls [21]. This model also enables cell-specific normalisation and was extended to detect differential expression between groups of cells [25]. Similar to the scDD simulation, Splatter provides a wrapper for the BASiCS simulation function, which is able to produce datasets with both endogenous and spike-in genes as well as multiple batches of cells. As the BASiCS simulation contains both biological and technical variation it can be used to test the ability of methods to distinguish between the two.

### Splat

We have developed the Splat simulation to capture many features observed in real scRNA-Seq data, including high expression outlier genes, differing sequencing depths (library sizes) between cells, trended gene-wise dispersion, and zero-inflation. Our model uses parametric distributions with hyper-parameters estimated from real data (Figure 1). The core of the Splat simulation is the gamma-Poisson hierarchical model where the mean expression level for each gene *i*, *i* = 1, …, *N*, is simulated from a gamma distribution and the count for each cell *j*, *j* = 1, …, *M*, is subsequently sampled from a Poisson distribution, with modifications to include expression outliers and to enforce a mean-variance trend.

**Figure 1:**
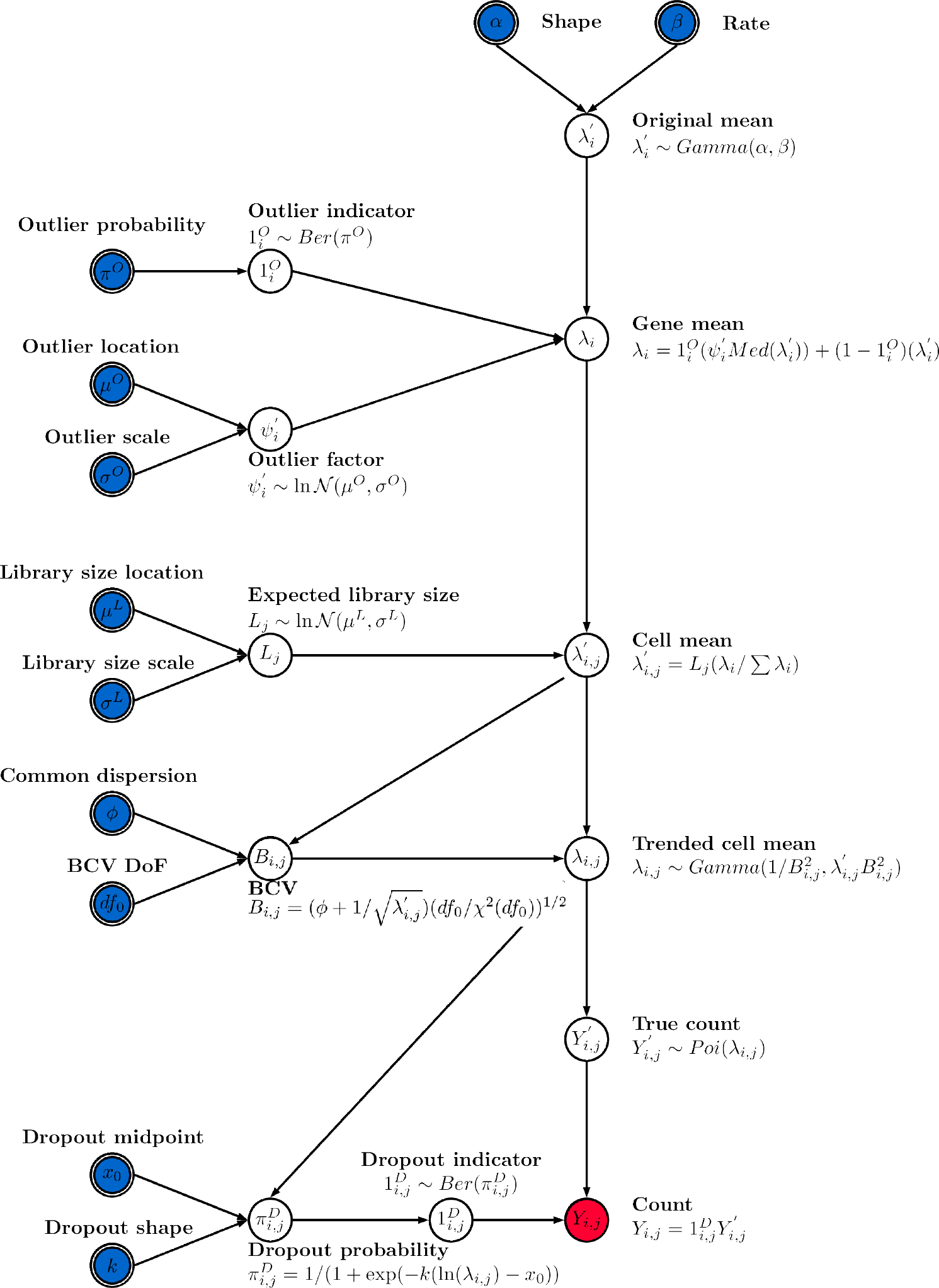
Diagram of the Splat simulation model. Input parameters are indicated with double borders and those that can be estimated from real data are shaded blue. Red shading indicates the final output. The simulation begins by generating means from a gamma distribution. Outlier expression genes are added by multiplying by a log-normal factor and the means are proportionally adjusted for each cell's library size. Adjusting the means using a simulated Biological Coefficient of Variation (BCV) enforces a mean-variance trend. These final means are used to generate counts from a Poisson distribution. In the final step dropout is (optionally) simulated by randomly setting some counts to zero, based on each gene's mean expression.

More specifically, the Splat simulation initially samples gene means from a Gamma distribution with shape α and rate *β*. While the gamma distribution is a good fit for gene means it does not always capture extreme expression levels. To counter this a probability (*π*^*0*^) that a gene is a high expression outlier can be specified. We then add these outliers to the simulation by replacing the previously simulated mean with the median of the simulated gene means multiplied by an inflation factor. The inflation factor is sampled from a log-normal distribution with location *μ^0^* and scale *σ^0^*.

The library size (total number of counts) varies within an scRNA-seq experiment and can be very different between experiments depending on the sequencing depth. We model library size using a log-normal distribution (with location *μ^L^* and scale *σ^L^*) and use the simulated library sizes (*L_j_*) to proportionally adjust the gene means for each cell. This allows us to alter the number of counts per cell independently of the underlying gene expression levels.

It is known that there is a strong mean-variance trend in RNA-Seq data, where lowly expressed genes are more variable and highly expressed genes are more consistent [26]. In the Splat simulation we enforce this trend by simulating the biological coefficient of variation (BCV) for each gene from a scaled inverse chi-squared distribution, where the scaling factor is a function of the gene mean. After simulating the BCV values we generate a new set of means (*λ_i,j_*) from a gamma distribution with shape and rate parameters dependent on the simulated BCVs and previous gene means. We then generate a matrix of counts by sampling from a Poisson distribution, with lambda equal to *λ_i,j_*. This process is similar to the simulation of bulk RNA-seq data used by Law et al. [27].

The high proportion of zeros is another key feature of scRNA-seq data [11], one cause of which is technical dropout. We use the relationship between the mean expression of a gene and the proportion of zero counts in that gene to model this process and use a logistic function to produce a probability that a count should be zero. The logistic function is defined by a midpoint parameter (*x*_0_), the expression level at which 50 percent of cells are zero, and a shape parameter (*k*) that controls how quickly the probabilities change from that point. The probability of a zero for each gene is then used to randomly replace some of the simulated counts with zeros using a Bernoulli distribution.

Each of the different steps in the Splat simulation outlined above are easily controlled by setting the appropriate parameters and can be turned off when they are not desirable or appropriate. The final result is a matrix of observed counts *Y_i,j_* where the rows are genes and the columns are cells. The full set of input parameters is shown in Table 1.

**Table 1:**
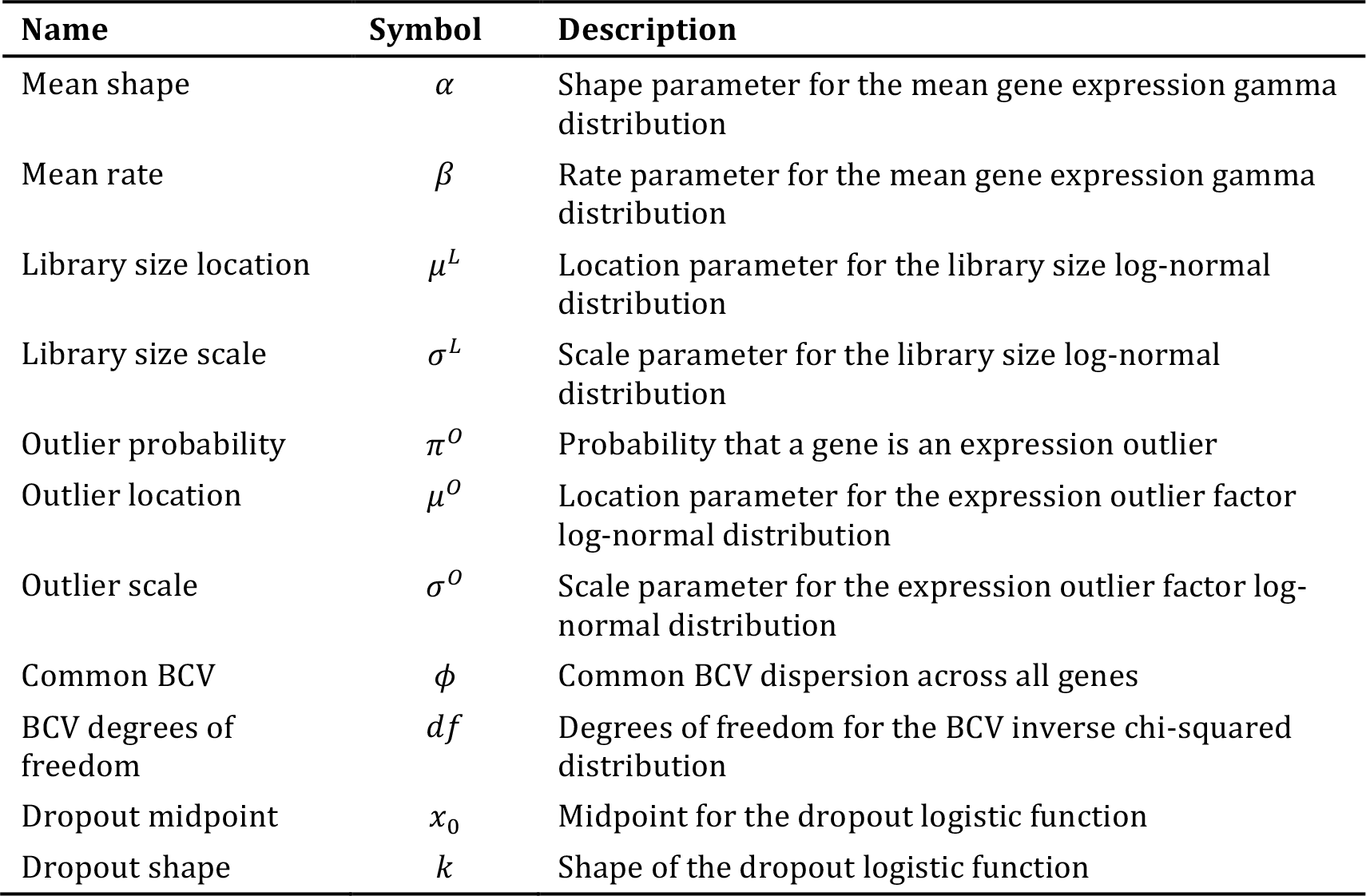
Input parameters for the Splat simulation model.

## Comparison of simulations

To compare the simulation models available in Splatter we estimated parameters from several real datasets and then generated synthetic datasets using those parameters. Both the standard and zero-inflated versions of the Splat and Lun 2 simulations were included, giving a total of eight simulations. We began with the Tung dataset which contains induced pluripotent stem cells from three HapMap individuals [28].

To reduce the computational time we randomly sampled 200 cells to use for the estimation step and each simulation consisted of 200 cells. Benchmarking showed a roughly linear relationship between the number of genes or cells and the processing time required (Additional File 1 Figures 4−5). The estimation procedures for the Lun 2 and BASiCS simulations are particularly time consuming, however the Lun 2 estimation can be run using multiple cores unlike the BASiCS estimation procedure. We did not perform any quality control of cells and only removed genes that were zero in all of the selected cells. We believe this presents the most challenging situation to simulate, as there are more likely to be violations of the underlying model. This scenario is also possibly the most useful as it allows any analysis method to be evaluated, from low-level filtering to complex downstream analysis. Figure 2 shows some of the plots produced by Splatter to compare simulations based on the Tung dataset.

**Figure 2:**
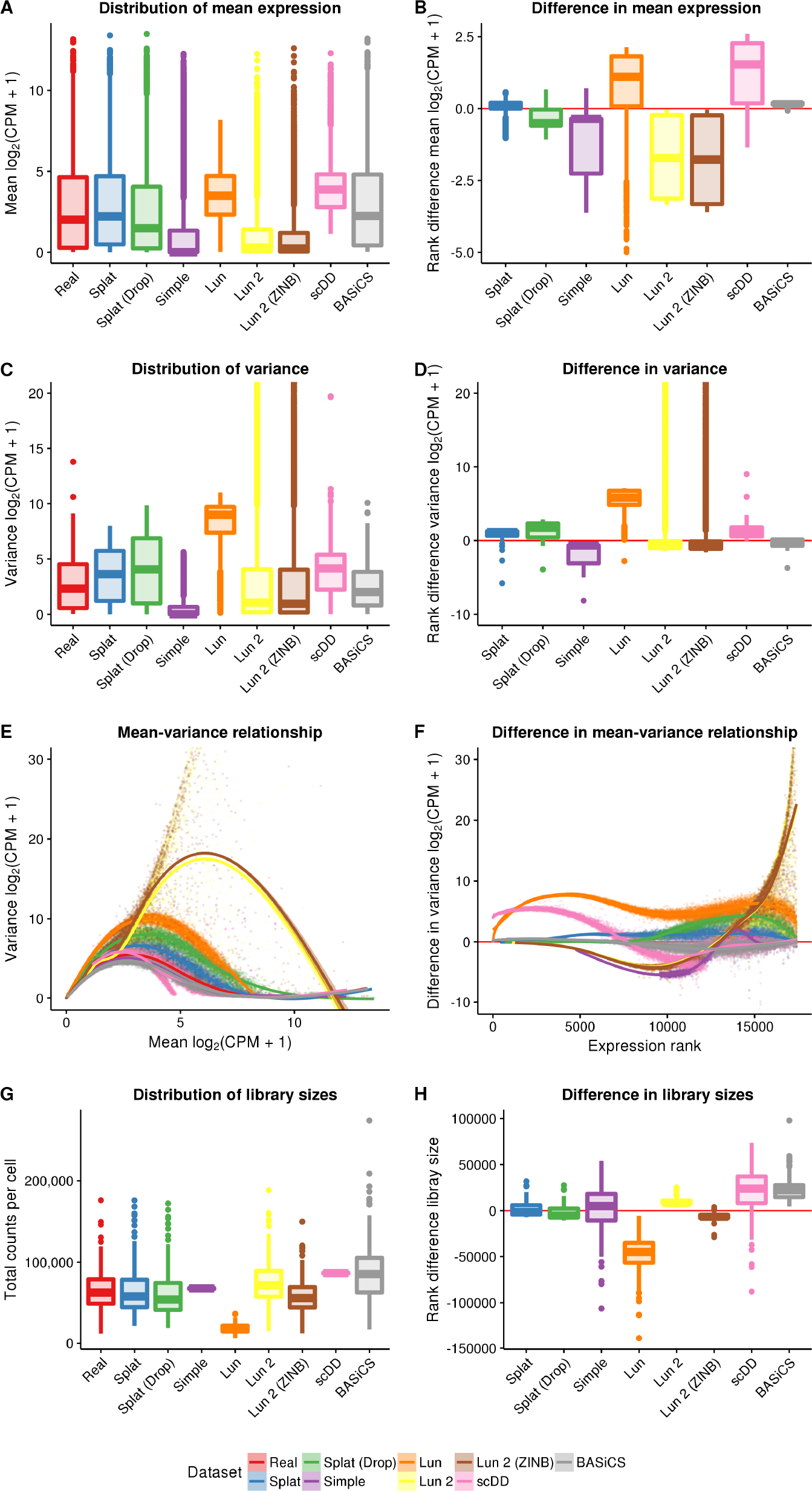
Comparison of simulations based on the Tung dataset. The left column panels show the distribution of mean expression (A), variance (C) and library size (G) across the real dataset and the simulations as boxplots, along with a scatter plot of the mean-variance relationship (E). The right column shows boxplots of the ranked differences between the real data and simulations for the same statistics: mean (B), variance (D), mean-variance relationship (F) and library size (H). Note that the y-axis for plots of the variance has been limited in order to show more detail. Variances for the Lun and Lun 2 simulations extend beyond what has been shown.

We compared the gene means, variances, library sizes and the mean-variance relationship. From these diagnostic plots, we can evaluate how well each simulation reproduces the real dataset and how it differs. One way to compare across the simulations is to look at the overall distributions (Figure 2, left column). Alternatively, we can choose a reference (in this case the real data) and look at departures from that data (Figure 2, right column). Examining the mean expression levels across genes, we see that the scDD simulation is missing lowly expressed genes, as expected, as is the Lun simulation. In contrast, the Simple and Lun 2 simulations are skewed towards lower expression levels (Figure 2A, Figure 2B). The BASiCS simulation is a good match to the real data as is the Splat simulation. Both versions of the Lun 2 simulation produce some extremely highly variable genes, an effect which is also seen to a lesser extent in the Lun simulation. The difference in variance is reflected in the mean-variance relationship where genes from the Lun 2 simulation are much too variable at high expression levels for this dataset. Library size is another aspect in which the simulations differ from the real data. The simulations that do not contain a library size component (Simple, Lun, scDD) have different median library sizes and much smaller spreads. In this example, the BASiCS simulation produces too many large library sizes, as does the Lun 2 simulation to a lesser degree.

A key aspect of scRNA-seq data is the number of observed zeros. To properly recreate an scRNA-seq dataset a simulation must produce the correct number of zeros but also have them appropriately distributed across both genes and cells. In addition, there is a clear relationship between the expression level of a gene and the number of observed zeros [29] and this should be reproduced in simulations. Figure 3 shows the distribution of zeros for the simulations based on the Tung dataset.

**Figure 3:**
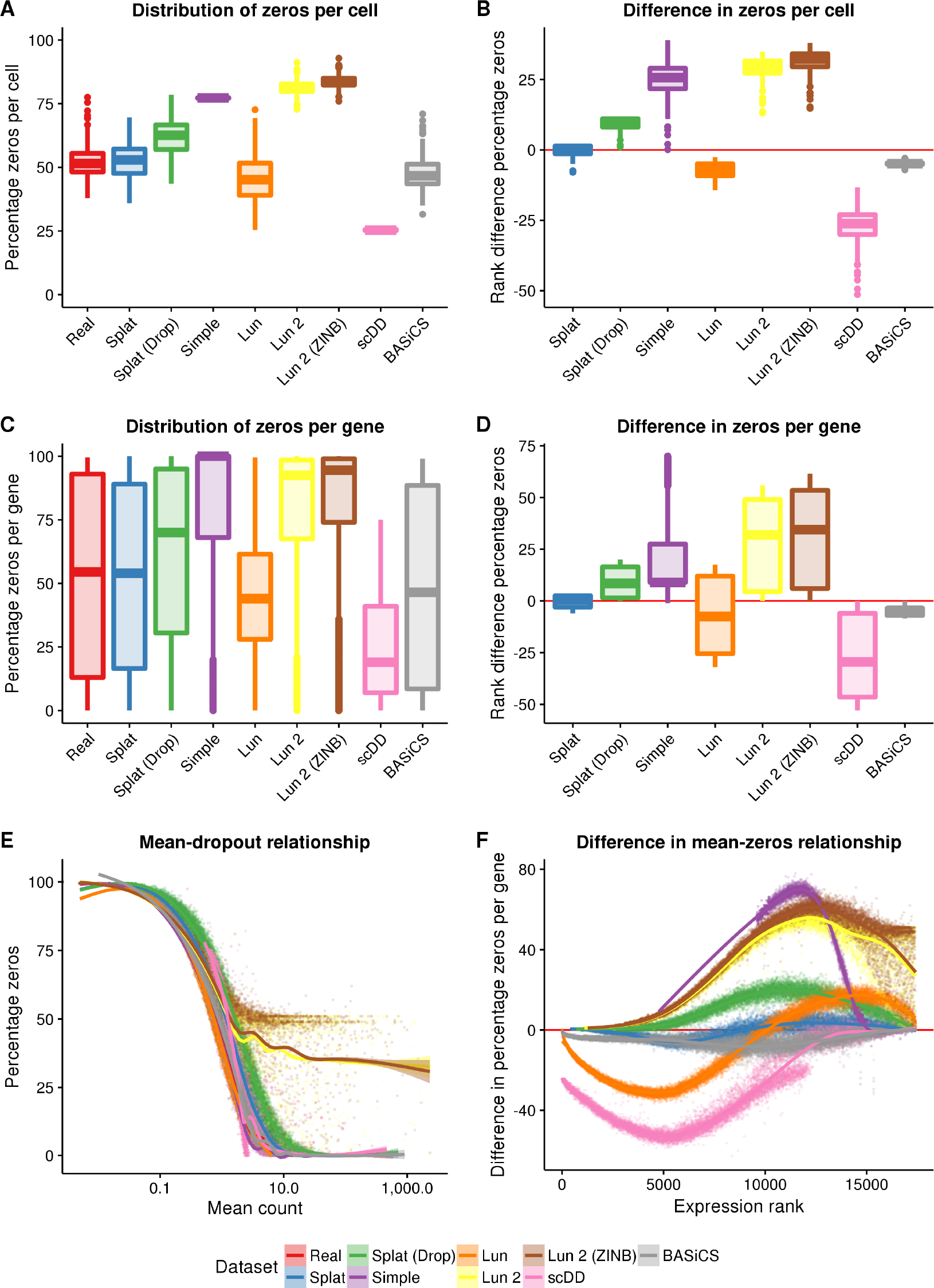
Comparison of zeros in simulations based on the Tung dataset. The top row shows boxplots of the distribution of zeros per cell (A) and the difference from the real data (B). The distribution (C) and difference (D) in zeros per gene are shown in the middle row. The bottom row shows scatter plots of the relationship between the mean expression of a gene (including cells with zero counts) and the percentage of zeros as both the raw observations (E) and as ranked differences from the real data (F).

For this dataset the Simple and Lun 2 simulations produce too many zeros across both genes and cells while the Lun and scDD simulations produce too few. Interestingly, the Splat simulation produces a better fit to this dataset when dropout is not included, suggesting that additional dropout is not present in the Tung dataset. However, this is not the case for all data and sometimes simulating additional dropout produces a better fit to the data (for example the Camp dataset presented below). We can also consider the relationship between the expression level of a gene, calculated including cells with zero counts, and the percentage of zero counts in that gene. The Lun and scDD simulations produce too few zeros at low expression levels, while the Simple and Lun 2 simulations produce too many zeros at high expression levels. It is important to note that as the scDD simulation removes genes with more than 75 percent zeros prior to simulation this model can never produce genes with high numbers of zeros as shown in Figure 3C. Both the Splat and BASiCS models are successful at distributing zeros across genes and cells as well as maintaining the mean-zeros relationship.

Although the analysis presented in Figure 2 and Figure 3 allows us to visually inspect how simulations compare with a single dataset we also wished to compare simulations across a variety of datasets. To address this we performed simulations based on five different datasets (outlined in Table 2) that varied in terms of library preparation protocol, cell capture platform, species and tissue complexity. Three of the datasets used Unique Molecular Identifiers (UMIs) [30] and two used full-length protocols. Complete comparison panels for all the datasets are provided in Additional File 1 Figures 5–10 and processing times for all datasets are shown in Additional File 1 Figure 11.

**Table 2:** Details of real datasets.

| Dataset | Species | Cell type | Platform | Protocol | UMI | Number of cells |
| --- | --- | --- | --- | --- | --- | --- |
| Camp [31] | Human | Whole brain organoids | Fluidigm C1 | SMARTer | No | 597 |
| Engel [32] | Mouse | Natural killer T cells | Flow cytometry | Modified Smart-seq2 | No | 203 |
| Klein [33] | Human | K562 cells | InDrop | CEL-Seq | Yes | 213 |
| Tung [28] | Human | Induced pluripotent stem cells | Fluidigm C1 | Modified SMARTer | Yes | 564 |
| Zeisel [34] | Mouse | Cortex and hippocampus cells | Fluidigm C1 | STRT-Seq | Yes | 3005 |

For each dataset, we estimated parameters and produced a synthetic dataset as described previously. We then compared simulations across metrics and datasets by calculating a median absolute deviation (MAD) for each metric. For example, to get a MAD for the gene expression means, the mean expression values for both the real data and the simulations were sorted and the real values were subtracted from the simulated values. The median of these absolute differences was taken as the final statistic. To compare between simulations, we ranked the MADs for each metric with a rank of one being most similar to the real data. Figure 4 summarises the ranked results for the five datasets as a heatmap. A heatmap of the MADs is presented in Additional File 1 Figure 12 and the values themselves in Additional File 2.

**Figure 4:**
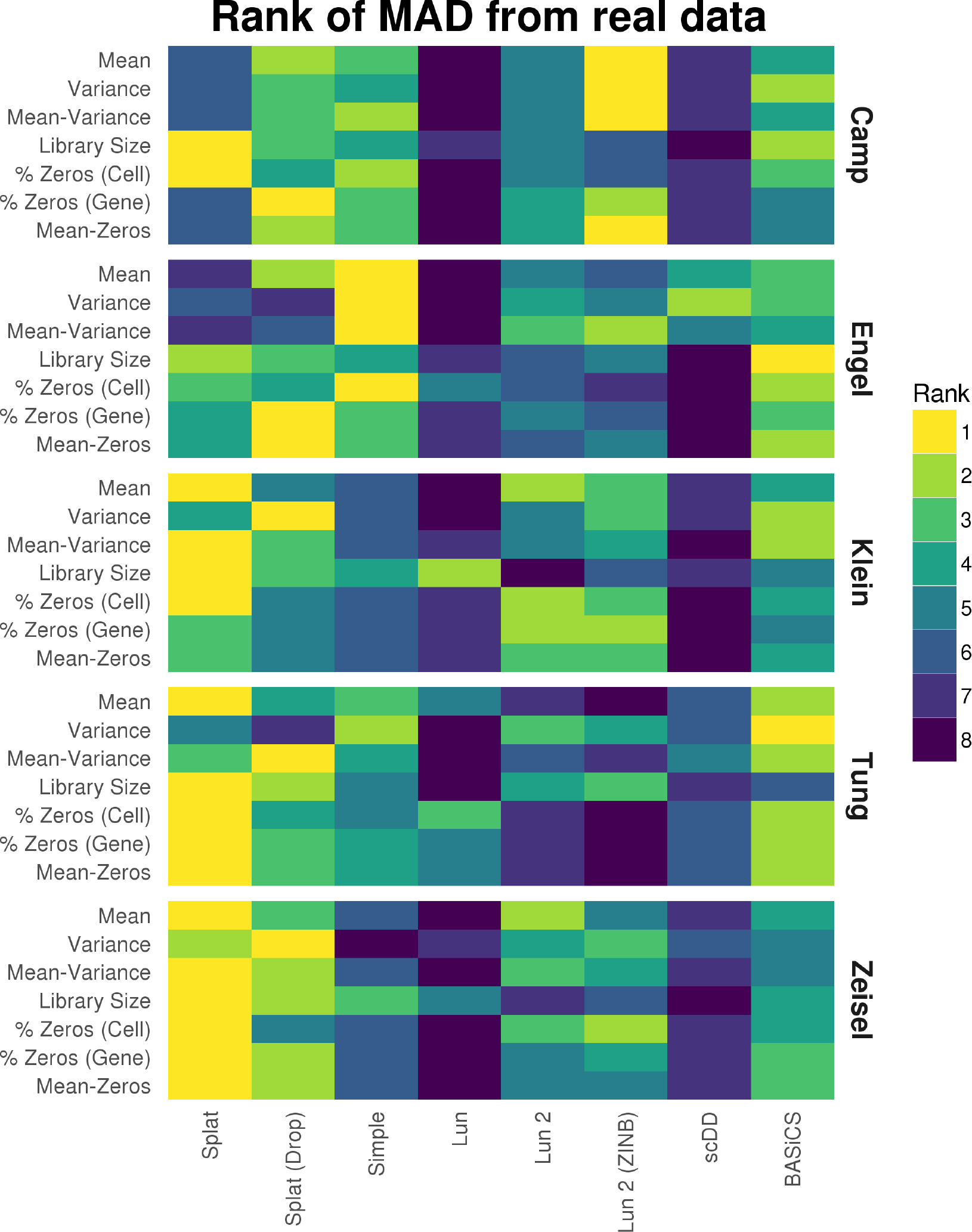
Comparison of simulation models based on various datasets. For each dataset parameters were estimated and synthetic datasets generated using various simulation methods. The Median Absolute Deviation (MAD) between each simulation and the real data was calculated for a range of metrics and the simulations ranked. A heatmap of the ranks across the metrics and datasets is presented here. We see that the Splat simulation (with and without dropout) performs consistently well, with the BASiCS simulation and the two versions of the Lun 2 simulation also performing well.

Looking across the metrics and datasets we see that the Splat simulations are consistently highly ranked. In general, it seems that the datasets are not zero-inflated and thus the zero-inflated simulations do not preform as well as their regular counterparts. The Splat simulations were least successful on the Camp cerebral organoid and Engel T-cell datasets. The complex nature of the Camp data (many cell types) and the full-length protocols used by both may have contributed to Splat's poorer performance. In this situation the semi-parametric, sampling-based models may have an advantage and the Lun 2 simulation was the best performer on most aspects of the Camp data. Interestingly, the Simple simulation was the best performer on the Engel dataset. This result suggests that the additional features of the more complex simulations may be unnecessary in this case or that other models may be more appropriate. The Lun simulation is consistently among the worst performing. However, given that this model is largely similar to the others, it is likely due to the lack of an estimation procedure for most parameters rather than significant problems with the model itself. The scDD simulation also often differed significantly from the real data, which is unsurprising as this simulation is designed to produce a filtered dataset, not the raw datasets used here. A comparison based on a filtered version of the Tung dataset, showing scDD to be a better match, is provided in Additional File 1 Figure 13.

Most importantly we see that simulations perform differently on different datasets. This emphasises the importance of evaluating different models and demonstrating their similarity to real datasets. Other comparisons may also be of interest for evaluation such as testing each simulated gene to see if it matches known distributions, an example of which is shown in Additional File 1 Figure 14. The Splatter framework makes these comparisons between simulation models straightforward, making it easier for researchers to choose simulations that best reflect the data they are trying to model.

## Complex simulations with Splat

The simulation models described above are sufficient for simulating a single, homogeneous population but not to reproduce the more complex situations seen in some real biological samples. For example, we might wish to simulate a population of cells from a complex tissue containing multiple mature cell types or a developmental scenario where cells are transitioning between cell types. In this section, we present how the Splat simulation can be extended to reproduce these complex sample types (Figure 5).

**Figure 5:**
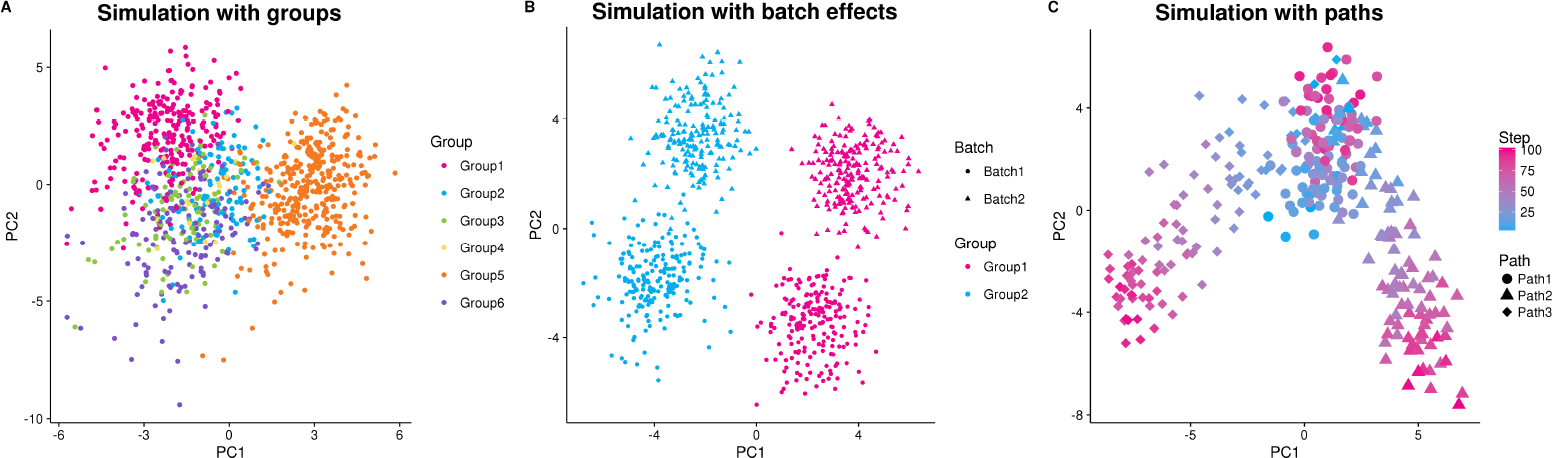
Examples of complex Splat simulations. (A) A Principle Components Analysis (PCA) plot of a simulation with six groups with varying numbers of cells and levels of differential expression. (B) A PCA plot of a simulation with two groups (pink and blue) and two batches (circle and triangle). PC1 separates groups (wanted biological variation) while PC2 separates batches (unwanted technical variation). (C) A PCA plot of a simulation with differentiation paths; the coloured gradient indicates how far along a path each cell is from blue to pink. A progenitor cell type (blue circles) differentiates into an intermediate cell type (pink circles/blue triangles or diamonds), which becomes one of two (pink triangle or diamond) mature cell types.

### Simulating groups

Splat can model samples with multiple cell types by creating distinct groups of cells where several genes are differentially expressed between the different groups. Previously published simulations can reproduce this situation to some degree but are often limited to fixed fold changes between only two groups. In the Splat simulation, however, differential expression is modeled using a process similar to that for creating expression outliers and can be used to simulate complex cell mixtures. Specifically a multiplicative differential expression factor is assigned to each gene and applied to the underlying mean. For DE genes, these factors are generated from a log-normal distribution while for other genes they are equal to one. Setting the number of groups and the probability that a cell comes from each group allows flexibility in how different groups are defined. Additionally, parameters controlling the probability that genes are differentially expressed as well as the magnitude and direction of DE factors can be set individually for each group. The resulting SCESet object contains information about which group each cell comes from as well as the factors applied to each gene in each group (Figure 5A).

### Simulating batches

A common technical problem in all sequencing experiments is batch effects, where technical variation is created during sample collection and preparation. The Splat simulation can model these effects using multiplicative factors that are applied to all genes for groups of cells. Adding this extra layer of variation allows researchers to evaluate how methods perform in the presence of unwanted variation (Figure 5B).

### Simulating paths

A common use of scRNA-seq is to study cellular development and differentiation. Instead of having groups of mature cells, individual cells are somewhere on a continuous differentiation path or lineage from one cell type to another. To model this, the Splat simulation uses the differential expression process described above to define the expression levels of a start and end cell for each path. A series of steps is then defined between the two cells types and the simulated cells are randomly assigned to one of these steps, receiving the mean expression levels at that point. Therefore, the simulation of lineages using Splat is defined by the differential expression parameters used to create the differences between the start and end of each path. It also incorporates the parameters that define the path itself, such as the length (number of steps) and skew (whether cells are more likely to come from the start or end of the path).

In real data it has been observed that expression of genes can change in more complex, non-linear ways across a differentiation trajectory. For example, a gene may be lowly expressed at the beginning of a process, highly expressed in the middle and lowly expressed at the end. Splat models these kinds of changes by generating a Brownian bridge (a random walk with fixed end points) between the two end cells of a path, which is then smoothed and interpolated using an Akima spline [35,36]. This random element allows many possible patterns of expression changes over the course of a path (Additional File 1 Figure 15). While non-linear changes are possible they are not the norm. Splat defines parameters that control the proportion of genes that are non-linear and how variable those genes can be.

Further complexity in simulating differentiation paths can be achieved by modeling lineages with multiple steps or branches. For example a stem cell that differentiates into an intermediate cell type that then changes into one of two mature cell types. These possibilities are enabled by allowing the user to set a starting point for each path (Figure 5C).

### Example: using Splatter simulations to evaluate a clustering method

To demonstrate how the simulations available in Splatter could be used to evaluate an analysis method we present an example of evaluating a clustering method. SC3 [5] is a consensus *k*-means based approach available from Bioconductor [37]. As well as assigning cells to groups, SC3 is able to detect genes that are differentially expressed between groups and marker genes that uniquely identify each group. To test SC3 we estimated Splat simulation parameters from the Tung dataset and simulated 400 cells from three groups with probabilities of 0.6, 0.25 and 0.15. The probability of a gene being differentially expressed in a group was 0.1, resulting in approximately 1700 DE genes per group. We then ran SC3 with three clusters (*k* = 3) and compared the results to the true groupings (Figure 6A). We also assessed the detection of DE and marker genes. True DE genes were taken as genes with simulated DE in any group and true marker genes as the subset of DE genes that were DE in only a single group (Figure 6B). This procedure was repeated 20 times with different random seeds to get some idea of the variability and robustness of the method.

**Figure 6:**
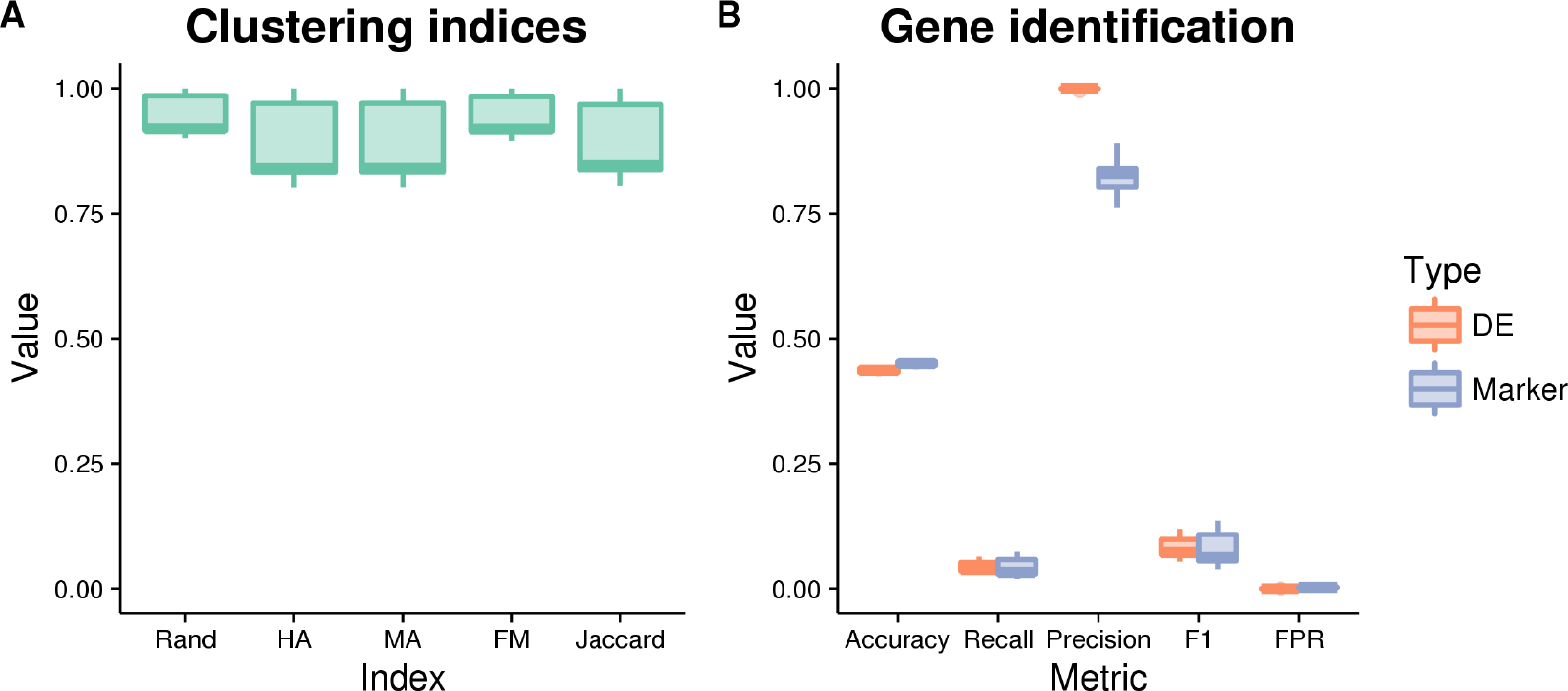
Evaluation of SC3 results. Metrics for the evaluation of clustering (A) include the Rand index, Hubert and Arabie's adjusted Rand index (HA), Morey and Agresti's adjusted Rand index (MA), Fowlkes and Mallows index (FM) and the Jaccard index. Detection of differentially expressed and marker genes were evaluated (B) using accuracy, recall (true positive rate), precision, F1 score (harmonic mean of precision and recall) and false positive rate (FPR). All of the metrics are presented here as boxplots across the 20 simulations.

Figure 6 shows the evaluation of SC3's clustering and gene identification on the simulated data. Five measures were used to evaluate the clustering: the Rand index (Rand), Hubert and Arabie's (HA) adjusted Rand index and Morey and Agresti's (MA) adjusted Rand index (both of which adjust for chance groupings), Fowlkes and Mallows index (FM) and the Jaccard index (Jaccard). All of these indices attempt to measure the similarity between two clusterings, in this case the clustering returned by SC3 and the true groups in the simulation. SC3 appears to identify clusters well for the majority of simulations, in some cases producing a near-perfect clustering. It may be interesting to examine individual cases further in order to identify when SC3 is able to perform better. Both the DE genes and marker genes identified by SC3 show a similar pattern across our classification metrics of accuracy, precision, recall and F1 score. On average approximately 2700 of the truly DE genes and 2500 of the true marker genes passed SC3's automatic filtering (with additional non-DE genes). SC3 then detected around 100 DE genes per simulation, along with 99 marker genes (median values). Precision (the proportion of identified genes that are true positives) is very high while recall (the proportion of true positives that were identified, or true positive rate) is very low. This tells us that in this scenario SC3 is producing many false negatives, but that the genes that it finds to be markers or DE are correct. This result is often desirable, particularly for marker genes, and is reflected in the very low false positive rate.

While it is beyond the scope of this paper, clearly this evaluation could be extended, for example by including more clustering methods, more variations in simulation parameters and investigating why particular results are seen. However, this data, and the code used to produce it, is an example of how such an evaluation could be conducted using the simulations available in Splatter.

## Discussion and conclusions

The recent development of single-cell RNA sequencing has spawned a plethora of analysis methods, and simulations can be a powerful tool for developing and evaluating them. Unfortunately, many current simulations of scRNA-seq data are poorly documented, not reproducible or fail to demonstrate similarity to real datasets. In addition, simulations created to evaluate a specific method can sometimes fall into the trap of having the same underlying assumptions as the method that they are trying to test. An independent, reproducible and flexible simulation framework is required in order for the scientific community to evaluate and develop sophisticated analysis methodologies.

Here we have developed Splatter, an independent framework for the reproducible simulation of scRNA-seq data. Splatter is available as an R package from Bioconductor and implements a series of simulation models. Splatter can easily estimate parameters for each model from real data, generate synthetic datasets and quickly create a series of diagnostic plots comparing different simulations and datasets.

As part of Splatter we introduce our own simulation called Splat. Splat builds on the gamma-Poisson (or negative binomial) distribution commonly used to represent RNA-seq data, and adds high-expression outlier genes, library size distributions, a mean-variance trend and the option of expression-based dropout. Extensions to Splat include the simulation of more complex scenarios, such as multiple groups of cells with differing sizes and levels of differential expression, experiments with several batches, or differentiation trajectories with multiple paths and branches, with genes that change in non-linear ways.

We performed an evaluation of the six simulation models currently available in Splatter by comparing synthetic data generated using estimated parameters to five published datasets. Overall Splat performed well, ranking highly on most metrics. However, other simulations performed better for some metrics or better reproduced specific datasets. We found the Camp cerebral organoid dataset the most challenging to simulate, perhaps because of the complex nature of this sample, which is comprised of many different cell types. In addition, this dataset (along with the Engel data) used a full-length protocol, which may contain additional noise compared to the UMI datasets [38].

One of the key features of scRNA-seq data is the high number of zero counts where no expression is observed for a particular gene in a particular cell. This can be especially challenging to simulate as not only must there be the correct number of zeros but they must be correctly distributed across genes and cells. We found that introducing dropout (in Splat) or zero-inflation (in Lun 2) often failed to improve the match to real datasets, suggesting that they are not truly zero-inflated. Together, the results demonstrate that no simulation can accurately reproduce all scRNA-seq datasets. They also emphasise the variability in scRNA-seq data, which arises from a complex set of biological (for example species, tissue type, cell type, treatment and cell cycle) and technical (for example platform, protocol, or processing) factors. Non-parametric simulations that permute real data could potentially produce more realistic synthetic datasets but at the cost of flexibility in what can be simulated and knowledge of the underlying parameters.

Finally, we demonstrated how Splatter could be used for the development and evaluation of analysis methods, using the SC3 clustering method as an example. Splatter's flexible framework allowed us to quickly generate multiple test datasets, based on parameters from real data. The information returned about the simulations gave us a truth to test against when evaluating the method. We found that SC3 accurately clustered cells and was precise in identifying DE and marker genes.

The simulations available in Splatter are well documented, reproducible and independent of any particular analysis method. Splatter's comparison functions also make it easy to demonstrate how similar simulations are to real datasets. Splatter provides a framework for simulation models, makes existing scRNA-seq simulations accessible to researchers and introduces Splat, a new scRNA-seq simulation model. As more simulation models become available, such as those replicating newer technologies including k-cell sequencing, they can be adapted to Splatter's framework. The Splat model will continue to be developed and may, in the future, include additional modules such as the ability to add gene lengths to differentiate between UMI and full-length data. We hope that Splatter empowers researchers to rapidly and rigorously develop new scRNA-seq analysis methods, ultimately leading to new discoveries in cell biology.

## Methods

### Splat parameter estimation

To easily generate a simulation that is similar to a given dataset, Splatter includes functions to estimate the parameters for each simulation from real datasets. Just as with the simulation models themselves, the estimation procedures are based on what has been published and there is variation in how many parameters can be estimated for each model. We have given significant attention to estimating the parameters for the Splat simulation. The parameters that control the mean expression of each gene (*α* and *β*) are estimated by fitting a gamma distribution to the winsorised means of the library size normalised counts using the fitdistrplus package [39]. The library size normalisation is a basic normalisation where the counts in the original dataset are adjusted so that each cell has the same number of total counts (in this case the median across all cells) and any genes that are all zero are removed. We found that genes with extreme means affect the fit of the gamma distribution and that this effect was mitigated by winsorising the top and bottom 10 percent of values to the 10th and 90th percentiles respectively. Parameters for the library size distribution (*μ^L^* and *σ^L^*) are estimated in a similar way by fitting a log-normal distribution to the unnormalised library sizes.

The procedure for estimating expression outlier parameters is more complex. Taking the library size normalised counts, outliers are defined as genes where the mean expression is more than two MADs greater than the median of the gene expression means. The outlier probability *π*^*0*^ is then calculated as the proportion of genes that are outliers. Parameters for the outlier factors (*μ*^*O*^ and *σ^O^*) are estimated by fitting a log-normal distribution to the ratio of the means of the outlier genes to the median of the gene expression means.

BCV parameters are estimated using the estimateDisp function in the edgeR package [22]. When testing the estimation procedure on simulated datasets we observed that the edgeR estimate of common dispersion was inflated (Additional File 1 Figure 16), therefore we apply a linear correction to this value (*ɸ̂* = 0.1 + 0.25 *ɸ̂*_edgeR_)

The midpoint (*x*_0_) and shape (*k*) parameters for the dropout function are estimated by fitting a logistic function to the relationship between the log means of the normalised counts and the proportion of samples that are zero for each gene (Additional File 1 Figure 17).

While we note that our estimation procedures are somewhat ad hoc, we found that these procedures are robust, efficient and guaranteed to produce parameter estimates on all datasets we tested.

### Datasets

Each of the real datasets used in the comparison of simulations is publicly available. Raw FASTQ files for the Camp dataset were downloaded from SRA (Accession SRP066834) and processed using a Bpipe (v0.9.9.3) [40] pipeline that examined the quality of reads using FastQC (v0.11.4), aligned the reads to the hg38 reference genome using STAR (v2.5.2a) [41] and counted reads overlapping genes in the Gencode V22 annotation using featureCounts (v1.5.0-p3) [42]. Matrices of gene by cell expression values for the Klein (Accession GSM1599500) and Zeisel (Accession GSE60361) datasets were downloaded from GEO. For the Tung dataset the matrix of molecules (UMIs) aligned to each gene available from https://github.com/jdblischak/singleCellSeq was used. This data is also available from GEO (Accession GSE77288). The Salmon [43] quantification files for the Engel dataset were download from the Conquer database (http://imlspenticton.uzh.ch:3838/conquer/) and converted to a gene by cell matrix using the tximport [44] package.

### Simulation comparison

For each dataset the data file was read into R (v3.4.0) [45] and converted to a gene by cell matrix. We randomly selected 200 cells without replacement and filtered out any genes that had zero expression in all cells or any missing values. The parameters for each simulation were estimated from the selected cells and a synthetic dataset generated with 200 cells and the same number of genes as the real data. Simulations were limited to 200 cells (the size of the smallest dataset) to reduce the computational time required. When estimating parameters for the Lun 2, scDD and BASiCS simulations cells were randomly assigned to two groups. For the Splat and Lun 2 simulations both the regular and zero-inflated variants were used to simulate data. The resulting eight simulations were then compared to the real data using Splatter's comparison functions and plots showing the overall comparison produced. To compare simulations across the datasets summary statistics were calculated. For each of the basic metrics (mean, variance, library size, zeros per gene and zeros per cell) the genes were sorted individually for each simulation and the difference from the sorted values and the real data calculated. When looking at the relationship between mean expression level and other metrics (variance, zeros per gene) genes in both the real and simulated data were sorted by mean expression and the difference between the metric of interest (eg. variance) calculated. The Median Absolute Deviation for each metric was then calculated and ranked for each dataset to give the rankings shown in Figure 4.

### Clustering evaluation

Parameters for Splat simulations used in the example evaluation of SC3 were estimated from the Tung dataset. Twenty synthetic datasets were generated using these parameters with different random seeds. Each simulation had three groups of different, with probabilities of 0.6, 0.25 and 0.1, and a probability of a gene being differentially expressed of 0.1. Factors for differentially expressed genes were generated from a log-normal distribution with location parameter equal to −0.1 and scale parameter equal to 0.3. For each simulation the SC3 package was used to cluster cells with *k* = 3 and asked to detect DE and marker genes, taking those with adjusted p-values less than 0.05. True DE genes were defined as genes where the simulated DE factor was not equal to one in one or more groups. Marker genes were defined as genes where the DE factor was not equal to one in a single group (and one in all others). Clustering metrics were calculated using the clues R package [46]. To evaluate the DE and marker gene detection we calculated the numbers of true negatives (TN), true positives (TP), false negatives (FN) and false positives. We then used these values to calculate the metrics shown in Figure 6: Accuracy (*Acc* = *TP* + *TN* / *Total number of genes*), Recall (*Rec* = *TP* / (*TP* + *FN*), Precision (*Pre* = *TP* / (*TP* + *FP*)), F1 Score (*F*1 = 2 ∗ (∗ *Rec* / (*Pre* + *Rec*))) and False Positive Rate (*FPR* = *FP* / (*FP* + *TN*)). Metrics were aggregated across the 20 simulations and boxplots produced using the ggplot2 package [47].

Session information describing the packages used in all analysis steps is included as Additional File 3. The code and dataset files are available at https://github.com/Oshlack/splatter-paper.

## Declarations

### Availability of data and materials

The datasets analysed during the current study are available from the repositories specified in the methods. The datasets and the code used to analyse them are also available from the repository for this paper https://github.com/Oshlack/splatter-paper (DOI: 10.5281/zenodo.833572). The Splatter package is available from Bioconductor (http://bioconductor.org/packages/splatter/) and is being developed on Github (https://github.com/Oshlack/splatter). The specific version of Splatter used in this paper is available at https://github.com/Oshlack/splatter/tree/v1.1.3-basics (DOI: DOI:10.5281/zenodo.833574).

## Competing interests

The authors declare no competing interests.

## Funding

Luke Zappia is supported by an Australian Government Research Training Program (RTP) Scholarship. Alicia Oshlack is supported through a National Health and Medical Research Council Career Development Fellowship APP1126157. MCRI is supported by the Victorian Government's Operational Infrastructure Support Program.

## Authors’ contributions

LZ developed the software and performed the analysis. BP contributed to the statistics and supervision. AO oversaw all aspects of the project. All authors contributed to drafting the manuscript.

## Acknowledgements

We would like to thank the authors of the BASiCS and scDD packages for their responses to our questions about how to include their simulations in Splatter as well as Mark Robinson and Charlotte Soneson for discussions regarding the simulation of scRNA-seq data. Our thanks also to Jovana Maksimovic and Sarah Blood for their comments on the manuscript.

## Additional files

**File name:** additional_figures_1-17.pdf

**File format:** PDF

**Title:** Additional figures

**Description:** Additional figures including: diagrams of other simulation models, Splatter comparison output for all datasets, example non-linear gene, dispersion estimate correction, mean-zeros fit, benchmarking and processing times.

**File name:** additional2_mads.csv

**File format:** CSV

**Title:** Table of MADs

**Description:** Table of the Median Absolute Deviations used to produce Figure 4 in CSV format.

**File name:** additional3_sessionInfo.pdf

**File format:** PDF

**Title:** Session information

**Description:** Details of the R environment and packages used for analysis.

